# Introduced species dominate different responses of grassland communities to climate change on serpentine and nonserpentine soils

**DOI:** 10.1101/844886

**Authors:** Joseph E Braasch, Maria A Johnson, Susan P Harrison, Katrina M Dlugosch

## Abstract

Introduced species are a common feature of modern plant communities and experience environmental challenges alongside native species. Changes to the environment may reveal distinct species-environment relationships for native and introduced components of plant communities. Extreme environmental change, such as drought, is predicted to result in declines in native species and increased opportunities for invasion, but empirical support for these ideas remains mixed. We tested for differences in the response of native and invaded species to environmental changes by analyzing a longterm dataset of species abundance in California grasslands collected during a period of severe drought. Sampling sites included a combination of stressful serpentine soils, which are resilient against invasion and maintain diverse native species assemblages, and more benign nonserpentine soils, which are heavily invaded and harbor low levels of native species cover. We found a significant correlation between sampling year and species composition for nonserpentine sites, but not for serpentine sites. These patterns were repeated when only introduced species were included in the analysis but no pattern of change was found for native species. The species most strongly associated with directional change on nonserpentine soils were three invasive Eurasian grasses, *Bromus hordaceus, Taeniatherium caput-medusae*, and *Avena fatua*. Differences in species composition on both serpentine and nonserpentine soils were significantly correlated with specific leaf area, a trait which has been linked to drought tolerance in these communities, although changes in abundance for the three Eurasian grasses most strongly associated with change did not consistently follow this pattern. Our analyses indicate relatively stable native community composition and strong directional change in introduced species composition, contradicting predictions for how native and introduced species will respond to environmental shifts, but supporting the hypothesis that native and invading species groups have important functional differences that shape their relationships to the environment.

## Introduction

Species introductions are ubiquitous features of modern biological communities, in some cases representing major components of biomass and species composition (Simberloff 1996; Bax et al. 2003; Powell et al. 2011; Vilà et al. 2011; Roy et al. 2016). Considerable efforts have been made to understand how introduced species establish and become invaders by investigating how they differ from the native species in their recipient communities (Rejmanek and Richardson 1996; Lee and Bell 1999; Sol et al. 2002; Van Kleunen et al. 2010). Explanations for the establishment of introduced species in novel habitats often focus on how species are able to occupy niche or functional space that is available either because native species have not evolved to exploit those resources, or because changing environments have opened additional niche space (Lyons and Schwartz 2001; Seabloom et al. 2003; Davis et al. 2011; Dlugosch et al. 2015). Explanations for invasion to high abundance emphasize that nonnative species might have greater competitive ability and the capacity to monopolize or more efficiently utilize limiting resources in the environment (Shea and Chesson 2002; MacDougall et al. 2009; van Kleunen et al. 2015). These hypotheses rely on the idea that native and nonnative species in a community have different functional ecologies, although evidence demonstrating such a distinction has been inconsistent and rarely neatly delineats functional differences between native and invading species (Thompson and Davis 2011; Kempel et al. 2013; Leffler et al. 2015).

Changes in the environment can be leveraged to better understand interspecific differences (Urban et al. 2012; Guisan et al. 2014) including those between native and introduced species. Climatic shifts have been shown to affect community composition (Cannone et al. 2007; Cantarel et al. 2013). diversity (Yang et al. 2011; Harrison et al. 2015). productivity (Arft et al. 1999: Rustad et al. 2001; Carlyle et al. 2014), and phenological or functional traits (Bradley et al. 1999; Walther 2010). These changes suggest that there are differences in the functional responses of individual species to environmental change, such that any functional differences between native and invading species are likely to result in divergent trajectories of these community components over time.

Climate is a ubiquitous force in shaping plant communities (Inouye 2008; Kimball et al. 2010; Wright et al. 2015), and anthropogenic climate change specifically is expected to make habitats more stressful, shuffling biological communities due to climatically driven range shifts, and causing local extirpation of native species (Parmesan 2006; Jump and Penuelas 2005; Menzel et al. 2006; Lurgi et al. 2012; Franklin et al. 2016). Climatic stress is expected to negatively impact the native component of biological communities either directly or from changes in species interactions (Van der Putten et al. 2010; Adler et al. 2012), food webs (Barton et al. 2009; Bannerman et al. 2011), and host-pathogen-vector relationships (Anderson et al. 2004; Garrett et al. 2006). The effects of environmental change on communities have been demonstrated through experimental manipulation (Philip Grime et al. 2000; Jägerbrand et al. 2009; Wolkovich et al. 2012) as well as through natural responses to climatic variation (Zheng et al. 2006; Elmendorf et al. 2015). Given that responses to climatic stress have been observed, long-term surveys of species abundances could be leveraged to better illuminate fundamental differences between the ecological roles of native and introduced species.

Perturbations to the environment are expected to increase opportunities for invasion when they promote disturbance, reduce the relative fitness or abundance of maladapted native species, and disrupt niche relationships (Shea and Chesson 2002; Wolkovich and Cleland 2011). Introduced species may be better able to respond to shifts in the environment if they possess greater adaptive capacity or plasticity (Molina-Montenegro et al. 2012; Chen et al. 2013; Wolkovich et al. 2013); properties which may also contribute to invasion (Bossdorf et al. 2005; Davidson et al. 2011). These ideas remain largely theoretical or are based on studies which focus on a small number of species responding to environmental treatments. Studies of long-term community dynamics could be particularly informative for testing both whether introduced species are functionally distinct, and specifically whether these species are uniquely benefiting from environmental change relative to native species.

In California, grassland ecosystems have been radically transformed through land use changes and subsequent invasion by exotic plants (Talbot et al. 1939; Murphy and Ehrlich 1989). This conversion began with the introduction of cattle and early 20th century farming practices to the region, which reduced native plant populations and introduced non-native weeds as cattle feed or forage contaminants in the mid 1800s (Menke 1989; Stromberg and Griffin 1996; Burcham 1956). Currently, introduced species dominate cover and biomass of these communities, although native species typically include similar species richness (Harrison et al. 2015). Many of the rare, intact native communities exist on stressful serpentine soils, which are distinct in their high heavy metal content, low water holding capacity, and lack of available nitrogen and phosphorus (Walker and B 1954; Kruckeberg 1984). Experiments have demonstrated that the stressful nature of these soils has contributed to their resilience against invasion (Huenneke et al. 1990; Harrison and Inouye 2002; Burge et al. 2016). The high native richness of these relict communities has also placed them as a conservation priority for the region (Murphy and Ehrlich 1989).

Environmental change has been identified as one of the major threats to the remnant native biodiversity in California grasslands (Harrison et al. 2008; Loarie et al. 2008; Damschen et al. 2010). Across much of the North American West, the severity of drought is predicted to increase in this already arid region (IPCC 2014; Griffin and Anchukaitis 2014; Robeson 2015; Swain et al. 2016). California recently experienced a decade long drought through the early 2000s, culminating in some of the lowest annual and seasonal precipitation on record from 2012 though 2014 (Copeland et al. 2016), similar to the long-term climate forecasts for this region. In California grasslands, water may act as a limiting resource, either directly or by affecting the availability of soil nutrients, with consequences for competitive outcomes (Seabloom et al. 2003; Enloe et al. 2004; Dukes et al. 2005; Everard et al. 2010). There is also evidence that drought has affected functional trait distributions of annual plant communities by favoring trait values that are expected to improve drought tolerance, such as lower specific leaf area (SLA) (Harrison et al. 2010; Hulshof et al. 2013; Eskelinen and Harrison 2015).

Here we test for divergent patterns of community change in native and invading plants in California grasslands during this recent period of extreme drought. We conduct a new analysis which leverages a long term plant community dataset collected by Harrison and colleagues (2015) in California from across co-occurring serpentine and nonserpentine grasslands. In a previous analysis of data from 2000-2014, Harrison et al. (2015) reported that species richness across both soil types declined, a pattern driven primarily by reductions in the abundance of rare, drought intolerant native forbs. The decline in species richness coincided with community-wide reductions specific leaf area (SLA), which is significantly correlated with winter precipitation and is consistent with community-wide response to drought (Harrison et al. 2015).

We use abundance data taken in this study from 2006-2016 to examine community compositional and functional change in a multivariate framework, and to identify any differential responses of native and introduced species to this period of extreme environmental conditions. Multivariate analyses offer the opportunity to examine shifts in the abundance of individual species responding to climate shifts community-wide, which might not be reflected in presence/absence changes that underlie trends in species richness. We use nonmetric multidimensional scaling (NMDS) to compare communities, splitting each community into its native and exotic components. This allowed us to describe the compositional change within each group independently of the other, because NMDS uses rank based abundances to calculate compositional differences. This is likely to be particularly important for detecting changes in the composition of native species on nonserpentine soils and introduced species on serpentine soils, which often represent minor components of their respective communities. We also use two separate tools to identify the individual species most strongly associated with directional change: gradient boosted models (GBM), a regression based machine learning tool, and principal component analysis (PCA) of the cumulative changes in species abundances over time.

We consider serpentine and nonserpentine communities separately given their distinct composition and ecology. We test for pairwise differences in community composition across years and for long-term, directional changes in community composition across multivariate space. We also assemble SLA trait values for the complete community, and test for directional change in this key functional trait. We ask whether native species dominate multivariate community change across soil types during drought, particularly through declines in abundance, in the same way that they dominated declines in species richness in previous analyses, and whether compositional change is associated with shifts to species with lower SLA, particularly non-native species.

## Methods

### Field Survey Data and Study Site

The long term data used for this study were collected by Harrison et al. (2015) and are reanalyzed here with the addition of new survey data for 2015 and 2016. The field data were collected at the University of California McLaughlin Reserve, California, USA (Napa and Lake Co.) from long term plots established in 2000. The region has a mediterranean-type climate, with the majority of annual rainfall occurring between the fall (October) and spring (April) seasons. Annual precipitation at the reserve has decreased in this region from 2003 to 2016, culminating in years with some of the lowest rainfall on record (Harrison et al. 2015a; Copeland et al. 2016). Annual plant communities within the reserve occupy stressful soils derived from serpentine rock and relatively fertile nonserpentine soils. Serpentine soils posses high levels of heavy metals (Mg and Ni), low levels of N and Ca, and low water holding capacity compared to nonserpentine soils (Proctor and Woodell 1975: Marrs and Proctor 1976: Kruckeberg 2006). Nonserpentine grasslands in California are heavily invaded by non-native grasses and forbs, primarily introduced from Eurasia (Jackson 1985: Mooney et al. 1986: Baker 1989). Serpentine soils are comparatively resilient against invasion (Huenneke et al. 1990: Going et al. 2009: Eskelinen and Harrison 2014) with low exotic species abundance. Previous work in these systems has shown that the plant communities found on the two soil types are distinct and respond differently to experimental manipulation of resources (Kruckeberg 1954: Whittaker 1954: Brady et al. 2005).

Long-term data were collected across 80 sites within the reserve (38 serpentine, 42 nonserpentine) spaced at least 50m apart. Each site was surveyed along a 40m transect with five permanent 1 m^2^ plots. From 2006 onward, species abundance was quantified with visual estimates of percent cover in each plot. Abundance prior to 2006 was recorded only as presence/absence and was not used in this study.

### Trait Data

Measurements of specific leaf area (SLA) (leaf area in mm^2^ / leaf dry mass in g) were collected by Harrison et al. (2015) for 111 species on serpentine and nonserpentine soils using protocols from Cornelissen et al. (2003). Many of the exotic species found at McLaughlin Reserve were not included in this survey, and SLA values for some species were only quantified on either serpentine (20) or nonserpentine (11) soils. When species lacked SLA measurements taken within McLaughlin Reserve, we either calculated a mean SLA value from all observations available in the Botanical Information and Ecology Network (BIEN) database or used SLA measurements from (Stevens et al. 2015), which were collected across a mixture of serpentine and nonserpentine sites in California using similar methods (Table SX). For species with SLA values measured on only one soil type, we used that SLA value for both soil types. There were 55 species for which no SLA value was available. Community weighted SLA values were calculated for each plot as the sum of each species’ SLA value multiplied by its percent cover. Site level SLA values were averaged over the 5 plots. A wilcoxon rank sum test was used to compare the distribution of SLA values for introduced and native species, with separate tests for serpentine and nonserpentine communities.

### Community change over time

For comparison to previous analyses (Harrison et al. 2015) which were conducted over a different timeframe (2000-2014), we tested whether changes in species richness from 2006 to 2016 were consistent with their previous observations. We used generalized linear models with species richness as the dependent variable, year as a fixed variable and site as a random variable with a poisson distribution using the ‘*lme4*’ package (Bates et al. 2014) in R version 3.5.3 (R Core Team 2013). We considered total species richness, native richness, and introduced species richness on serpentine and nonserpentine sites separately.

For abundance analyses, cover data for each site was averaged across all 5 plots for further analyses. Compositional differences between sites for 2006 to 2016 were evaluated with non-metric multidimensional scaling, hereafter ‘NMDS’ (Field et al. 1982; Kruskal 1964b; Shepard 1962) with the *‘vegan’* package v2.5-4 in R (Dixon 2003). Ordinations were constrained to two dimensions for ease of visualization and use in downstream analyses, and were conducted using Bray-Curtis distances with weighted average scores.

The quality of an NMDS ordination is most often represented as its stress, the cumulative standardized differences between the original distance matrix and the matrix representing differences in ordination space modeled via a stepwise, monotonically increasing function (Kruskal 1964a). Stress increases with the number of observations in the analysis, however, and may not be appropriate for evaluating the quality of NMDS ordinations of large datasets, such as those with over 100 observations (Clarke 1993; Dexter, et. al 2018), as we have in this study. Instead, we compared NMDS output to permutation-based ecological null communities as suggested by Dexter et al. (2018). Null communities were produced by randomly shuffling species observations to produce a novel distance matrix using the ‘*oecosimu*’ function from the *‘vegan’* package in R, with the *‘swsh_swap’* permutation method (Dixon 2003). Stress values from the original dataset were then compared to the distribution of stresses for 99 null communities with a z-distribution. Significant results from this test indicate that NMDS is able to depict an underlying ecological pattern within the real data, which is not present in null communities (Dexter et al., 2018). We performed these analyses separately for serpentine and non-serpentine communities, and for exotic and native plants each within serpentine soils and nonserpentine soils (6 analyses in total).

To visualize differences between years, ellipses were plotted representing 30% confidence intervals for each year. To test for directional changes in community composition across time, and in traits, we fit explanatory vectors for year and weighted SLA onto all significant community NMDS plots with the function envfit in the vegan package (Dixon 2003). To test for any effect of year on community composition, and for the duration of time required to detect this difference, we used a permutation based analysis of variance (PERMANOVA function from the *‘vegan’* package) to compare the pairwise difference in species abundances from two different years to 999 pairs of randomly shuffled communities. This was repeated for all pairwise year to year comparisons for serpentine and nonserpentine communities.

### Identification of species most responsible for community differences

Individual species contributions to community change cannot be extracted from NMDS axes, which are non-linear, and so we used a machine learning approach to identify which species best predicted community change in NMDS ordination space. We used Gradient Boosted Models (GBM) to predict NMDS centroid values for each year and along the two NMDS axes. Generally, GBMs are a class of machine learning tool which produce a prediction model, in this case predicting NMDS axis values for yearly centroids from species cover data, in the form of decision trees, similar to random forest models. The predictive power of the decision tree is optimized by producing a secondary loss function, which describes the difference between predicted and observed values, and then by using derivatives to find parameter values which minimize the loss function. To test the predictive ability of GBM outputs, the dataset was randomly split into a training set (⅔ of observations per year) and testing set (⅓ of observations per year) prior to running the GBM. The ability of the GBM to accurately predict NMDS axis values with the testing data was calculated as the mean squared error, with better runs possessing lower error. We ran each GBM simulation 1000 times, because the optimization algorithm has the potential to get stuck at local minima. To run the GBM we used function *‘gbm’* from the R package gbm v2.1.5 (Ridgeway et al. 2006). Each model run included 4500 trees (n.trees = 4500), with only one way interactions (interaction.depth =1), a minimum of three observations on terminal nodes (n.minobsinnode = 3), and a learning rate of 0.1 (shrinkage = 0.1). Parameters were optimized by using the ‘train’ function from the *‘caret*’ package (Kuhn 2008).

We also performed a principal component analysis on the cumulative change in species abundances over time, using the function *‘prcomp’* in R (R Core Team 2013). We extracted the top 5 species loadings for the first two PC axes and compared these with the GBM predictions.

## Results

The ten years of observation between 2006 and 2016 included 239 species, the majority of which were native (eventual reference for dataset location). On serpentine soils there were 190 species (148 native, 42 exotic), and on nonserpentine soils there were 173 species recorded (128 native, 51 introduced). Harrison et al. observed species richness declines on both serpentine and nonserpentine soils from 2000-2014, and we verified that these patterns did not change in our dataset spanning 2006 to 2016 (Table S1, Figure S1). Reductions in the richness were significant for both native and exotic species on both serpentine and nonserpentine soil (Table S1). Species richness for native species declined at a faster rate than introduced species on nonserpentine soils (regression slope for native spp.= −0.354, for introduced spp. = −0.18; Figure S1). On serpentine soils, native species richness declined more slowly than introduced species richness (regression slope for native spp. = 0.20, for introduced spp. = 0.33; Figure S1).

In both serpentine and nonserpentine habitats, we found significant differences between communities across years (PERMANOVA: serpentine: F=6.6, P < 0.001; nonserpentine: F=30.8, P < 0.001). Moreover, the majority of all possible pairwise comparisons of composition between years were significantly different (nonserpentine: 44/55 comparisons; serpentine: 38/55 comparisons; Figure S2). Comparisons of community composition across consecutive years were significant for 70% of all tests on nonserpentine soil and 50% of all tests on serpentine soils.

Ordination of nonserpentine sites had significantly lower stress than null communities (*z*=-41.1, P<0.01; Figure S3). When vectors for year and site-level SLA were fit in NMDS space, both showed significant correlations with community composition (year: r^2^=0.102, P < 0.001; SLA: r^2^=0.075, P < 0.001; Figure 1B). Additionally, the vectors for year and SLA were oriented at an obtuse angle (□=159.3°) implying a negative relationship between them in NMDS space. Ordination of native species on nonserpentine soils was not better than null communities (*z*=1.0, P=0.31; Figure 2A, Figure S3) while ordination for exotic species was significantly better than null (*z*=-21.2, P<0.01; Figure 2B, Figure S3). Additionally, there was no significant correlation between differences in NMDS space and year (r^2^=0.0112, P=0.077), or SLA (r^2^=0.0025, P=0.543). Similar to results for the entire community, both year and weighted-SLA were significant when mapped onto NMDS space for exotic, nonserpentine plants (year: r^2^=0.148, P < 0.001; SLA: r^2^=0.075, P < 0.001; Figure 2B), with a negative relationship between them (□=137.9°).

**Figure 1.**
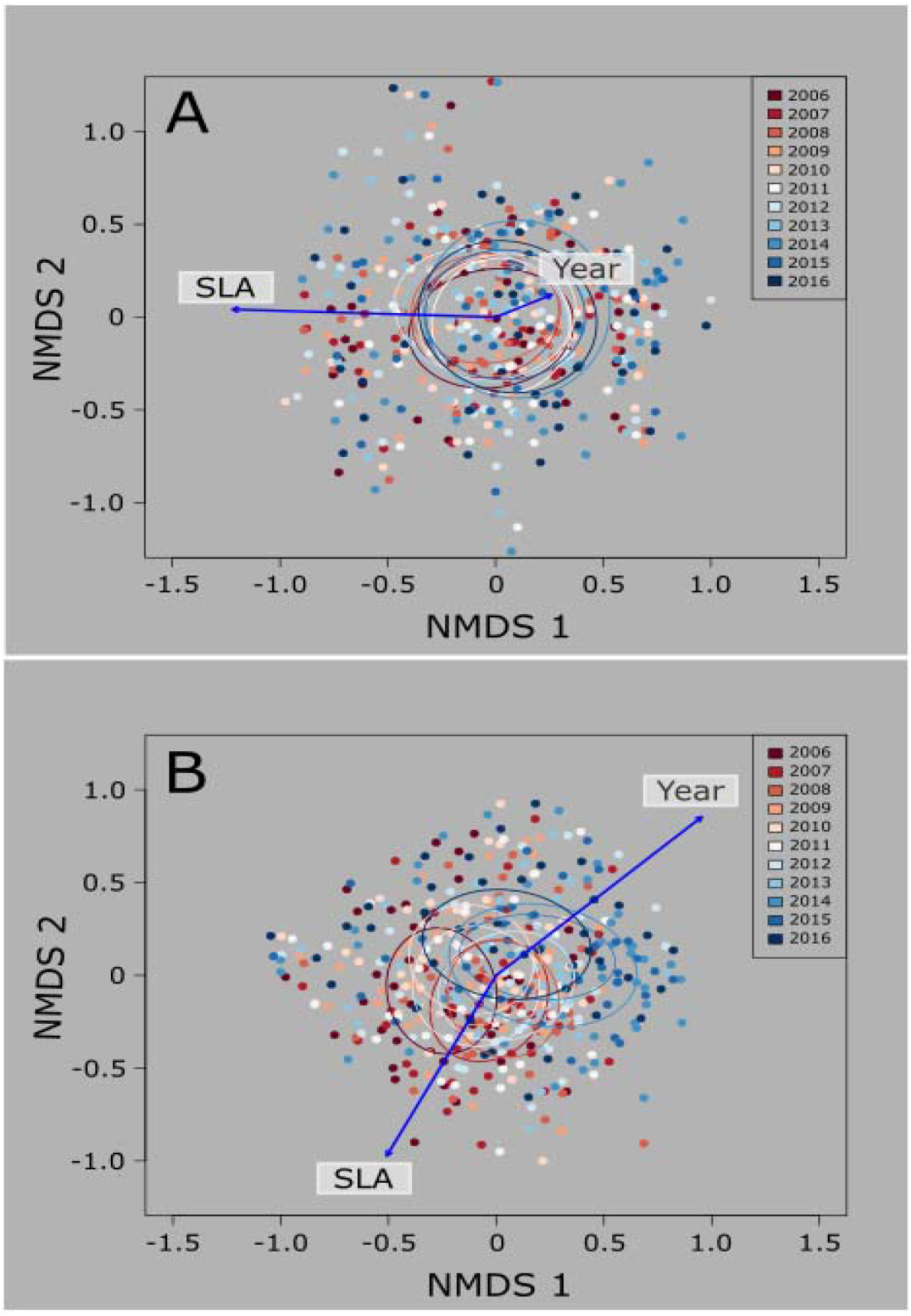
Nonmetric multidimensional scaling ordinations in species space from 2006 to 2016 with fitted vectors for year and community weighted specific leaf area (blue arrows). Individual sites (dots) and 30% confidence ellipses for each year are colored by year. On serpentine sites (A), differences in site-level species composition are significantly correlated with community weighted specific leaf area (P < 0.001), but not the year in which the survey was performed (P = 0.32). In nonserpentine sites, both year and community weighted specific leaf area were significantly correlated with compositional differences (P < 0.001 for both effects), and point in opposite directions, which indicates that year and specific leaf area may be negatively correlated in ordinated space.

**Figure 2.**
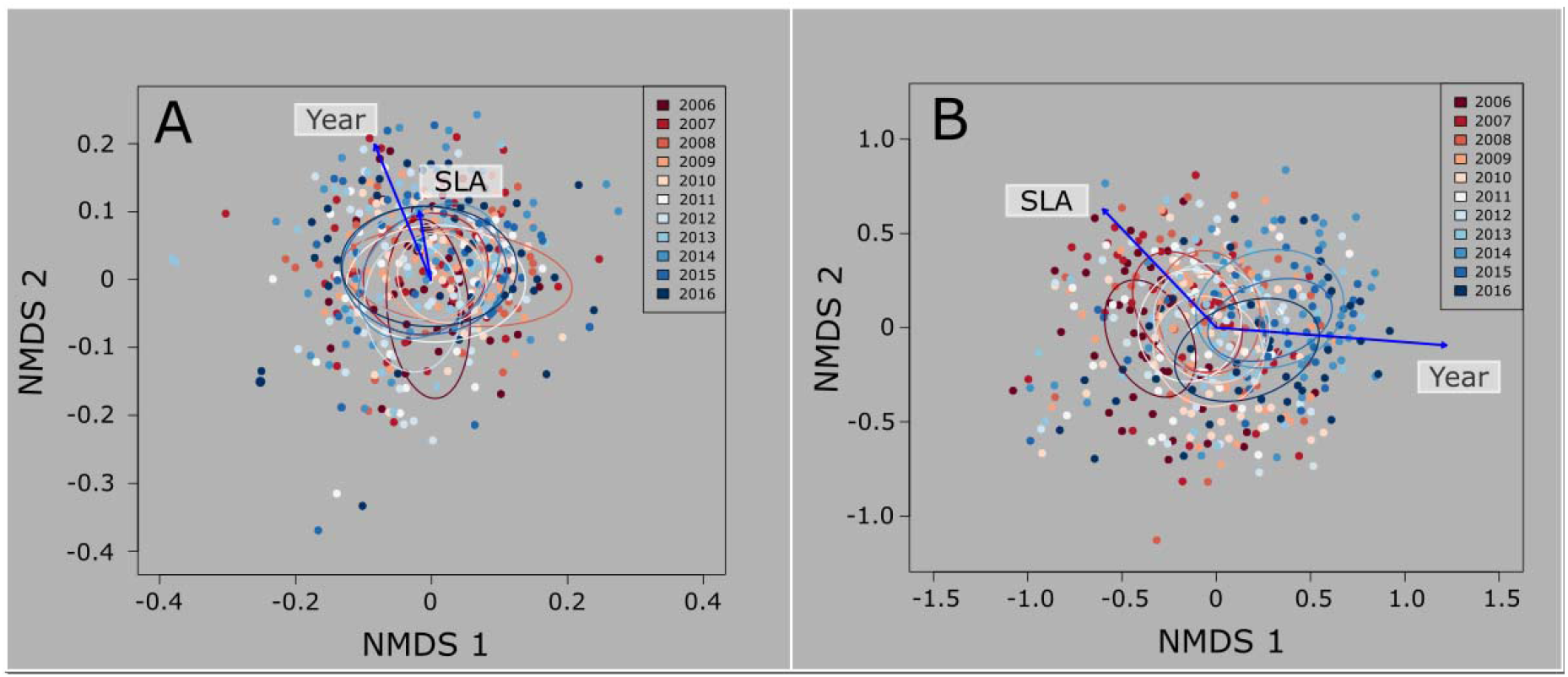
Nonmetric multidimensional scaling ordinations in species space for native (A) and exotic (B) species on nonserpentine soils from 2006 to 2016 with fitted vectors for year and community weighted specific leaf area (blue arrows). Individual sites (dots) and 30% confidence ellipses for each year are colored by year. Differences in site-level species composition are not significantly correlated with community weighted specific leaf area or year for native species (A) (Year: P=0.077; SLA: P=0.542), but are both significantly correlated for exocic species (B) (Year: P<0.001; SLA: P<0.001). The relationship between SLA and year was negative for exotic species.

NMDS ordination for the serpentine community from 2006 to 2016 yielded significantly lower stress than null communities (stress = 0.24, *z*=-66.6, P<0.01; Figure S3), supporting the presence of underlying ecological structure. When we fit vectors onto NMDS space, there was no significant directional correlation of community structure with year (r^2^=0.005, P = 0.34), while there was a significant correlation with site-level SLA (r^2^=0.093, P < 0.001; Figure 1A). Results were similar when only the native component of the serpentine community was analyzed; NMDS ordination had lower stress than null communities (stress = 0.25, *z*=-47.8, P<0.01; Figure S3) and fitted vectors for year and SLA were non-significant and significant respectively (Year: r^2^=0.007, P = 0.262; SLA: r^2^=0.121, P < 0.001; Figure 2A).

For exotic species on serpentine soil, NMDS ordination did not produce a significantly better fit than for null communities (stress = 0.22, *z*= −.45, P= 0.43; Figure S3), but this may be due to the unusual structure of the species matrix. Although stress for this ordination was lower than the majority of null communities, random permutation of the data yielded 15 out of 99 runs with lower stress, including one run with stress very close to zero, which could result from the outsized influence of a small number of observations on the ability of the NMDS algorithm to reach find the best possible solution. Performing the ordination in three dimensions did not resolve this behavior, and the algorithm failed to converge in more than half of all attempted ordinations due to the composition of a small number of sites around the periphery of the ordination space. Although NMDS ordination did not perform better than null communities, fitted vectors for year and community-weighted SLA were both significant (year: r^2^=0.071, P < 0.001; SLA: r^2^=0.051, P < 0.001; Figure 3B). Year and SLA were also positively correlated in NMDS space with an angle of □=44.4° between them.

**Figure 3.**
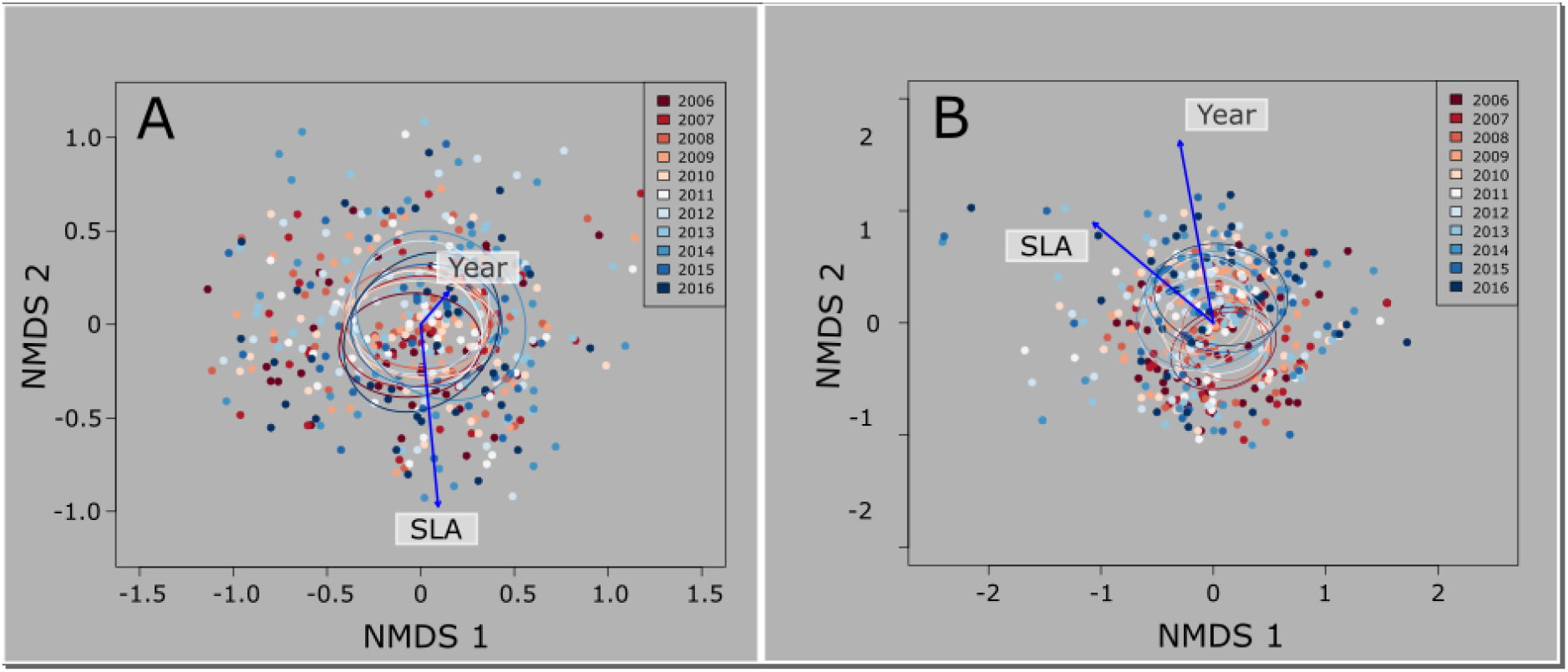
Nonmetric multidimensional scaling ordinations in species space for native (A) and exotic (B) species on serpentine soils from 2006 to 2016 with fitted vectors for year and community weighted specific leaf area (blue arrows). Individual sites (dots) and 30% confidence ellipses for each year are colored by year. For native species (A), differences in site-level species composition are significantly correlated with community weighted specific leaf area (P < 0.001), but not the year in which the survey was performed (P = 0.26). Year and community weighted specific leaf area were significantly correlated with compositional differences in exotic species (P < 0.001 for both effects). The vectors for these effects point in similar directions, indicating a positive relationship between them in ordination space.

To identify the species most responsible for community change in the nonserpentine community we used gradient boosted models to predict changes along NMDS axes from a subset of species abundance data. Model mean squared error for predictions along NMDS axis 1 ranged from 0.077 to 0.101 with an average of 0.088, and mean squared error along NMDS axis 2 ranged from 0.065 to 0.085 with an average of 0.074. GBM models consistently assigned the greatest weights for predictive ability to a set of common exotic and invading species. Along NMDS axis 1, the highest weighted species for all 1000 GBM models was *Bromus hordaceus*, followed by *Taeniatherium caput-medusae*, and *Avena fatua* (Figure 4A), all of which are annual Eurasian grasses common to invaded western landscapes. These three grasses were also the most important species for making accurate predictions along NMDS axis 2, but with less difference among them (Figure 4B). Native species were rarely ranked in the top five species for accurate predictions along either NMDS axis, with two native species identified as infrequent rank 4 and 5 predictors for NMDS axis 1 (*Clarkia purpurea* and *Dichelostemma capitatum)* and axis 2 *(Holocarpha virgata* and *Lupinus bicolor*).

**Figure 4.**
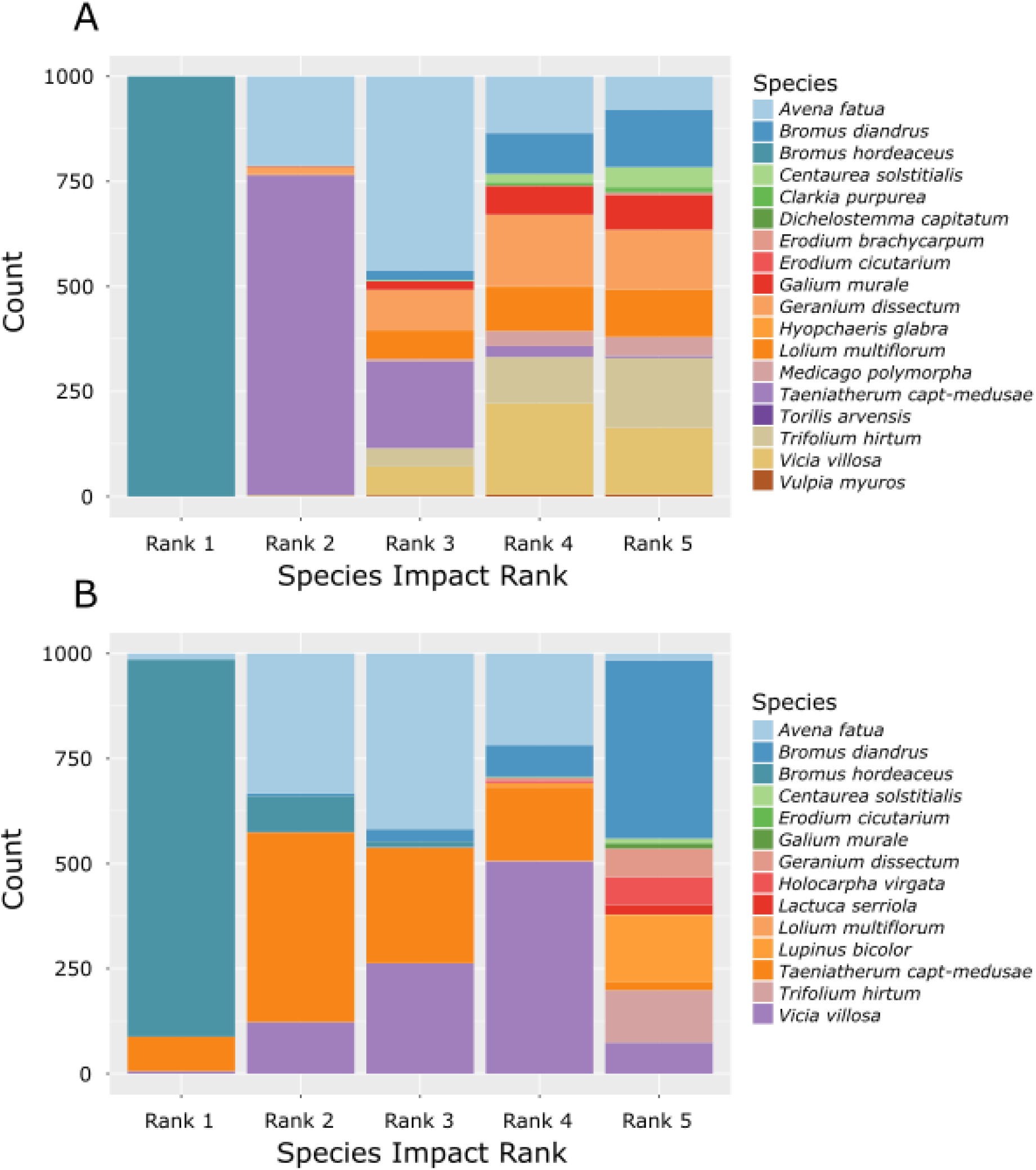
Results from 1000 gradient boosted models which predicts NMDS year centroids in figure X from individual nonserpentine site-level observations. Species are ranked by their order of importance for producing accurate predictions. A) predictions along NMDS axis 1; B) predictions along NMDS axis 2.

We conducted PCA ordination on the cumulative differences in species abundances over time. Axis loadings for PC axis 1 and 2 were largest for the same three introduced grasses identified by GBM analysis; *B. hordaceous, T. caput-medusae*, and *A. fatua* (Figure 5) The set of five species with the greatest loading along the first and second PC axes did not include native species. Two of these species, *B. hordaceus* and *T. caput-medusae*, decreased in abundance over the survey period, while *A. fatua* increased (Figure S5). The SLA for all three grasses was above the mean SLA for plants in this community (*A. fatua*: 314 mm^2^/g; *B. hordaceous*: 311 mm^2^/g; *T. caput-medusae*: 245 mm^2^/g; mean: 245 mm^2^/g; Figure S5). Overall, introduced species on both nonserpentine and serpentine soils had significantly greater SLA than native plants (nonserpentine: W=4715, P<0.001; serpentine: W=4964, P<0.001) (Figure S4).

**Figure 5.**
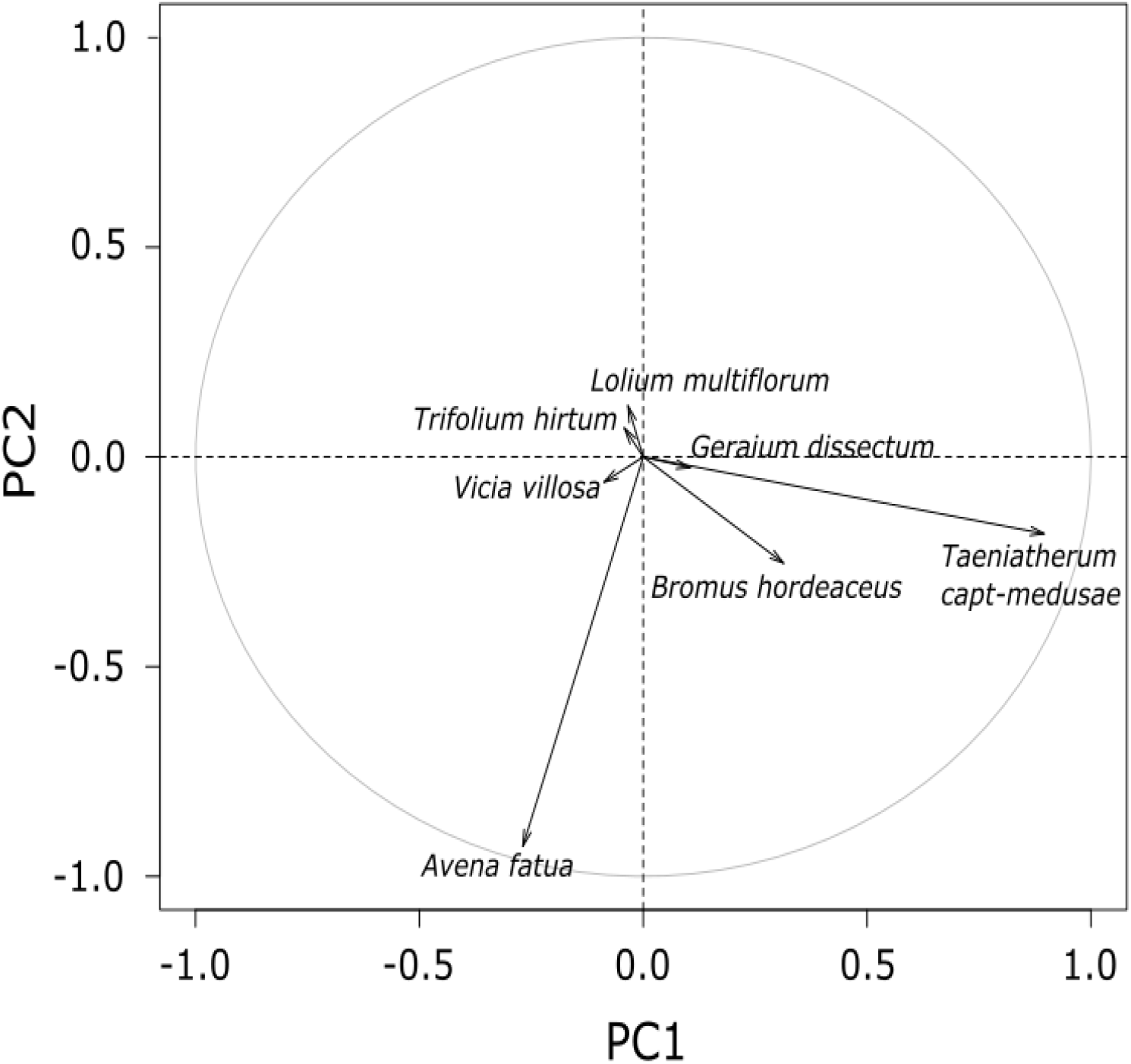
Axis loadings from principal coordinates analysis of changes in species composition over time for nonserpentine sites. Only the top five species loadings for each axis are shown.

## Discussion

Introduced species are now common features of plant communities, but their functional differences from native species (if any) remains a poorly resolved question in ecology. By analyzing a long term plant community dataset, we find that directional change in community composition occuring during a period of unprecedented drought is primarily attributable to shifting abundances of introduced species. These changes were correlated with reductions in SLA, a functional trait which is negatively correlated with performance under drought conditions (Harrison et al. 2010; Sandel et al. 2010; Dwyer et al. 2014). Corresponding patterns of directional change were not observed in the native component of these plant communities, despite a significant association between SLA and differences in native community composition. Our results point to unique functional differences between native and introduced species which may only be evident in the presence of long-term environmental stress.

Water is a limiting resource in California grasslands, and the years of this study had some of the lowest total annual rainfall on record (Copeland et al. 2016), which has been associated with reduced abundance of drought intolerant species and communities with lower weighted SLA (Harrison et al. 2015). Our analysis of nonserpentine sites also captured this relationship between community change and SLA. Directional change was negatively correlated with weighted SLA in NMDS space on nonserpentine soil when we considered the entire community or only introduced species. The abundant, introduced grasses in this system are known to compete strongly for water which affects their growth and fecundity (Suttle et al. 2007). Water addition has also been shown to negate the competitive effect of introduced grasses on native grass growth and fecundity (Hamilton et al. 1999). Native species as a whole did not undergo directional change on nonserpentine sites, which could be explained by their already lower SLA values and greater drought tolerance. A similar study of community change at these sites found greater variability in species richness in response to annual differences in precipitation on heavily invaded nonserpentine communities than on serpentine communities and that greater community stability was linked to communities with more drought tolerant species (Fernandez-Going et al. 2012). Our results support the idea that introduced species may generally be constrained to functional niche space that is more novel for the community and more likely to be lost or gained under environmental stress and variation (Goergen and Daehler 2002; Diez et al. 2012; Wolkovich and Cleland 2014).

Gradient boosted models and PCA of cumulative changes in species abundance provided further evidence that introduced species, and invading annual grasses specifically, are most strongly associated with directional change. We identified the grasses *A. fatua, B. hordaceus*, and *T. caput-medusae* as the most influential species for producing accurate GBM predictions and the species with the greatest axis loadings in a PCA of cumulative change in species abundance. *Avena fatua* quadrupled in abundance by the end of the survey (not including an outlier year in 2015 when its cover increased by an order of magnitude), while *B. hordaceus*, and *T. caput-medusae* became less abundant. We expected the species most strongly associated with directional change to increase or decrease based on their SLA values in accordance with community wide patterns. Of these three grasses only the high SLA, *T. caput-medusae* matched the community wide pattern by decreasing over the course of the study. The other grasses have almost identical SLA values that are close to the community median, but these species exhibited differing population trajectories. Other traits might be useful for explaining the changes in cover for these two species. Phenology traits, for example, can act as alternative mechanisms for coping with drought though drought escape and the efficacy of specific drought tolerance strategies may also change with the competitive environment (Fotelli et al. 2001; Maestre et al. 2009).

SLA was significantly related to compositional differences for the NDMS ordinations of serpentine plants which performed better than null communities (all plants and native plants only). There was no directional change in these sites, so SLA based differences are likely a product of persistent microhabitat variration. The ‘harshness’ of serpentine soils can vary with consequences for plant community traits and may have greater effects than variation in annual precipitation (Johnston and Proctor 1981; Wright et al. 2006; Eskelinen and Harrison 2014). This underlying structure in soil composition could mask signals of community change in the context of our analyses, where we did not track the behavior of individual sites. It is also possible that plants growing on these soils are less strongly affected by drought due to adaptation for serpentine’s naturally low water holding capacity. Increasing water availability does not significantly impact serpentine plant communities without nutrient supplementation, and the addition of both water and nitrogen decreases the abundance of low SLA species on these soils (Eskelinen and Harrison 2015). Studies conducted on infertile limestone soils have also found that these plant communities are resilient to warming and drought due to a suite of traits which facilitate persistence in environments with scarce resources. (Philip Grime et al. 2000; Grime et al. 2008). Our results contribute to this small body of work suggesting that plant communities on stressful soils may be preadapted to some environmental shifts and extremes (Philip Grime et al. 2000; Tielbörger et al. 2014; Harrison et al. 2015b).

We had predicted that directional change would be more strongly associated with native species, given previous work finding that reductions in species richness were the result of the loss of rare native forbs across serpentine and nonserpentine communities and the high variability in abundance of native forb cover (Fernandez-Going et al. 2012; Harrison et al. 2015). Our inclusion of additional sampling from 2015 and 2016 did not change patterns of species loss on nonserpentine soils. Surprisingly, we found differing patterns of compositional change across these two communities which appears to be related to the relative dominance of introduced species. The discrepancy between patterns of compositional change and richness loss may result from how individual species are weighted in NMDS analysis. Considerations of richness weigh each species equally, regardless of abundance.

NMDS ordination performed with Bray-Curtis distances reduces the influence of common species by using rank order abundance, but rare species will still have relatively small individual effects in highly diverse communities such as those considered here. On heavily invaded nonserpentine soils, introduced species comprise the majority of plant cover and biomass while native species often represent ~0.1% of plant cover and occur infrequently across sampling sites. The drop out of rare native species from individual plots will therefore have small effects on our analyses of community composition, which will be dominated by shifts in abundant species. The introduced species found on nonserpentine soils are also common across all sampling sites (e.g. there is low □-diversity), which may also increase our ability to detect signals of change in this subset of the community relative to native species on nonserpentine or serpentine communities with higher □-diversity.

Our analyses of community composition do suggest that abundant native and introduced plants occupy different functional niches resulting in compositionally stable native communities and unstable introduced communities in the presence of environmental extremes. Theoretical models of how native and introduced species will respond to environmental change assume change in native communities will create opportunities for invasion (Shea and Chesson 2002; Wolkovich and Cleland 2011). but native species within this system are compositionally stable. Greater functional diversity has been shown to confer resistance to invasion in plant communities (Pokorny et al. 2005; Hooper and Dukes 2010), and rare or uncommon species can have disproportionate importance in producing these effects (Zavaleta and Hulvey 2004; Hulvey and Zavaleta 2012). In our analyses, the loss of rare native species is not associated with overall increases in the abundance of introduced species. Rather, introduced species appear to trade-off with one another in abundance in response to extreme drought while native communities remained compositionally stable. Similar relationships between native and introduced species were observed in European plant communities where native species dynamics were stable and changed in accordance with landscape scale environmental variation, whereas patterns of change in newly arrived exotic species were more stochastic and associated with short term climatic variation (Latombe et al. 2018). Fluctuating abundance of introduced species do not appear to result in the recovery of native populations as a result, at least on short timescales, and instead allow for other introduced species to become more compositionaly dominant.

Compositional changes during a period of exceptional drought could be particularly informative for understanding plant community responses to long term climate change. Rare but extreme events can exhibit some of the strongest selection on populations inducing phenotypic change that far outlasts the duration of the selective event (Brown and Brown 1998; Bailey and van de Pol 2016; Grant et al. 2017) with significant consequences for community composition (Thibault and Brown 2008; Boucek and Rehage 2014). Introduced species will prove particularly useful for studying the process of climate adaptation if they possess unique functional traits combinations which have been lost to selection in native species through a history of extreme selective events.

Functional differences between native and introduced species have been central to the theory of biological invasions dating back to Darwin (Darwin 1859) with empirical support from experimental microcosm studies (Zavaleta and Hulvey 2004; Hulvey and Zavaleta 2012). but less consistent support from the field (Meiners 2007; Moles et al. 2012; van Kleunen et al. 2015). We found that extreme drought in California grasslands revealed functional differences between native and introduced species. In particular, drought revealed trade-offs in abundance among dominant introduced species, and stability among common native species, suggesting that extreme environmental events are favoring certain functional types but not invading species as a whole. These patterns were only apparent in association with long term responses to environmental change, which may be essential for observing consistent differences between native and invader function.

## Funding

USDA ELI Fellowship #2017-67011-26034 to J.B.

USDA grant #2015-67013-23000 and NSF #1750280 to K.M.D

Any support for Maria (Dukes?)

Susan please add

---

**Table S1.** Changes in species richness for different plant communities estimated using generalized linear mixed models.

| Soil |  | estimate | z-value | P-value |
| --- | --- | --- | --- | --- |
| Nonserpentine |  |  |  |  |
|  | All | -0.024 | -7.6 | <0.001 |
|  | Native | -0.033 | -7.09 | <0.001 |
|  | Introduce | -0.017 | -3.94 | <0.001 |
| Serpentine |  |  |  |  |
|  | All | -0.019 | -6.68 | <0.001 |
|  | Native | -0.016 | -4.98 | <0.001 |
|  | Introduce | -0.034 | -5.143 | <0.001 |

**Figure S1.**
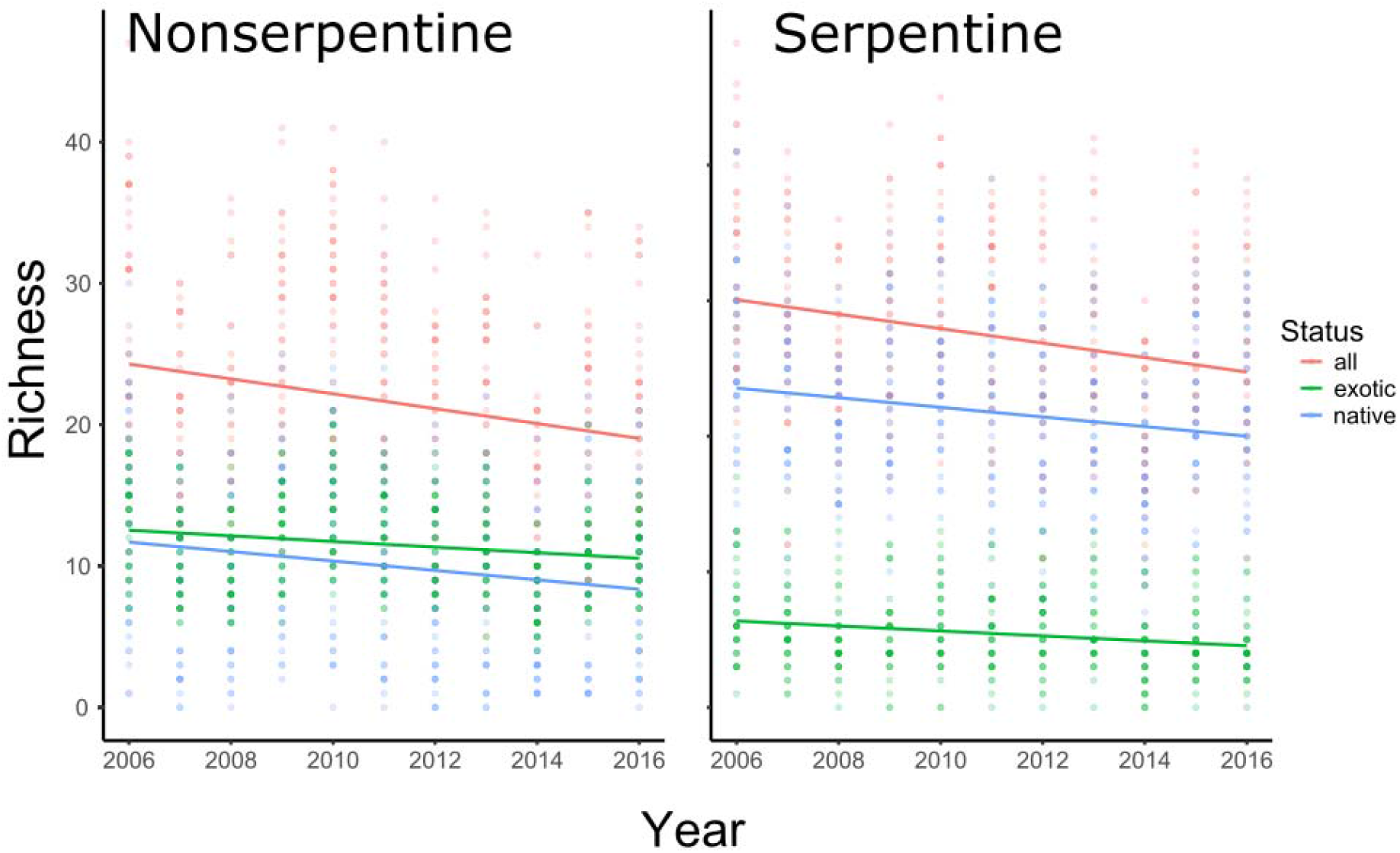
Changes in species richness on nonserpentine and serpentine soils.

**Figure S2.**
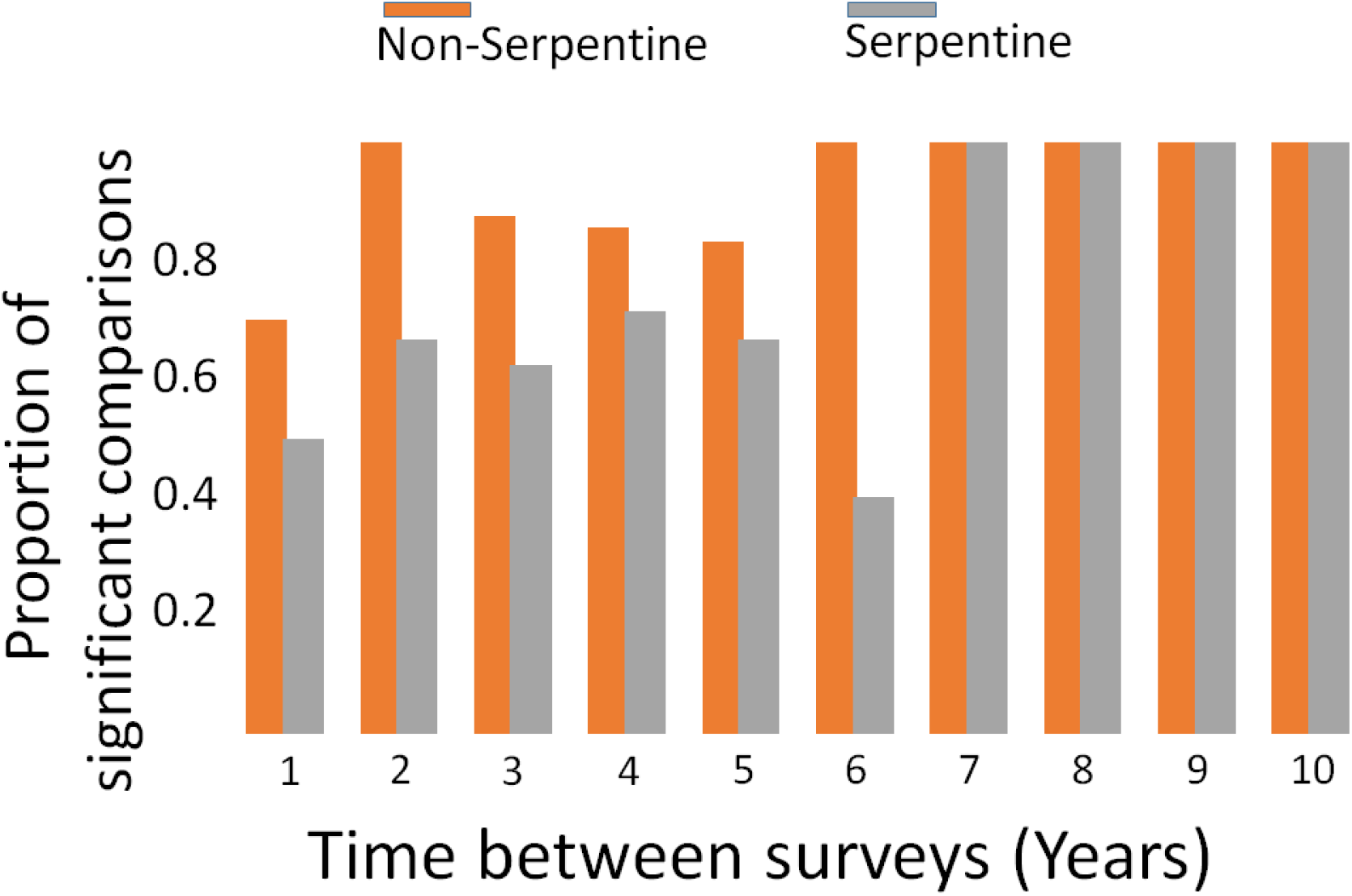
Proportion of significant (P < 0.05) pairwise community comparisons across all possible pairs of years between 2006 and 2016 years. For example, for each community there are ten 1-year difference comparisons, and one 10-year comparison. Orange = Nonserpentine sites, Grey = Serpentine sites. Pairwise comparisons were performed using permutation based analysis of variance (PERMANOVA).

**Figure S3.**
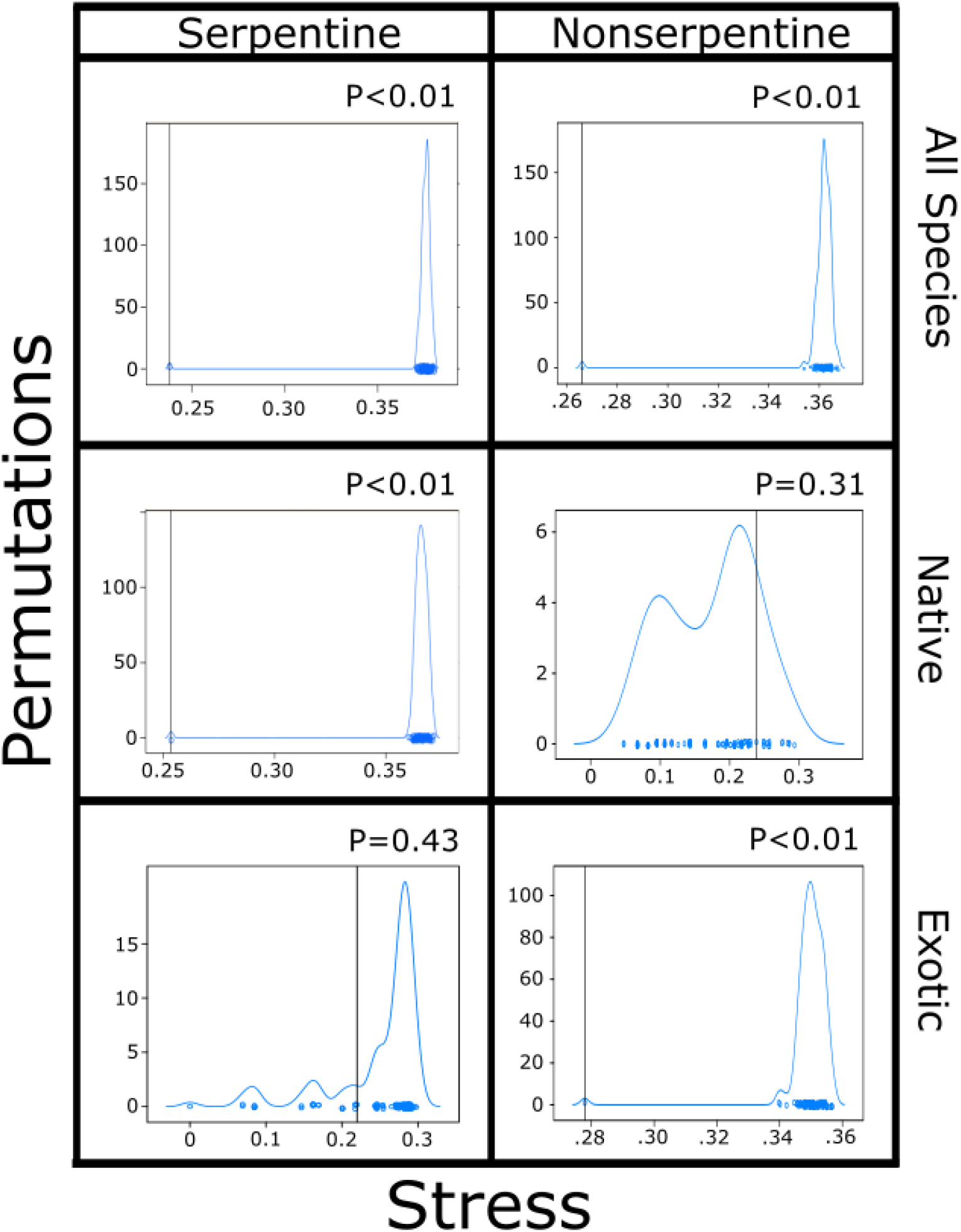
Distribution of NMDS ordination for null communities on serpentine and nonserpentine soils. Ordination values for for observed communities are depicted as vertical lines. P-values are calculated from z-scores.

**Figure S4.**
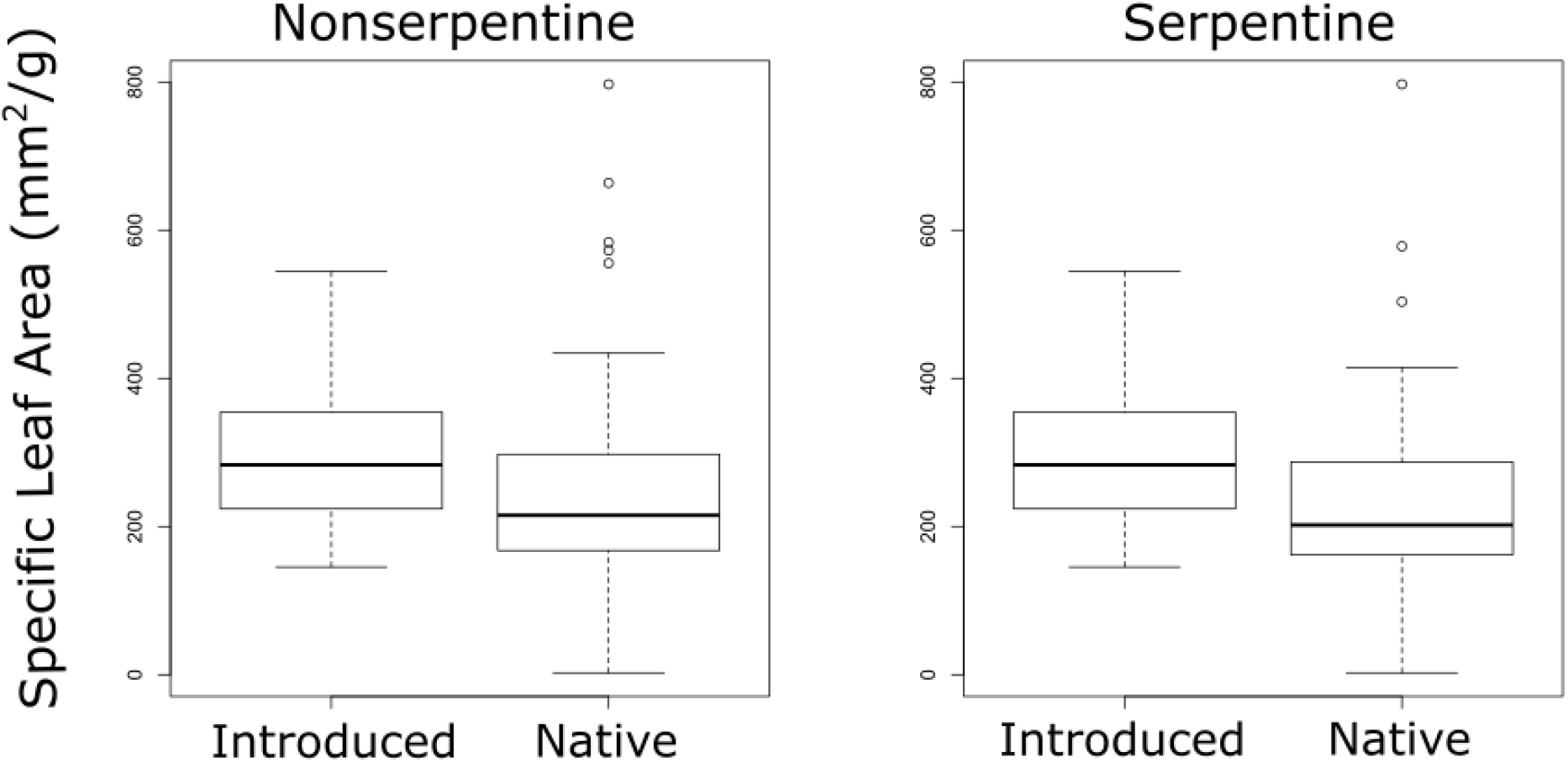
The distribution of specific leaf area for introduced species is significantly greater (P<0.001) than for native species on both nonserpentine and serpentine soils.

**Figure S5.**
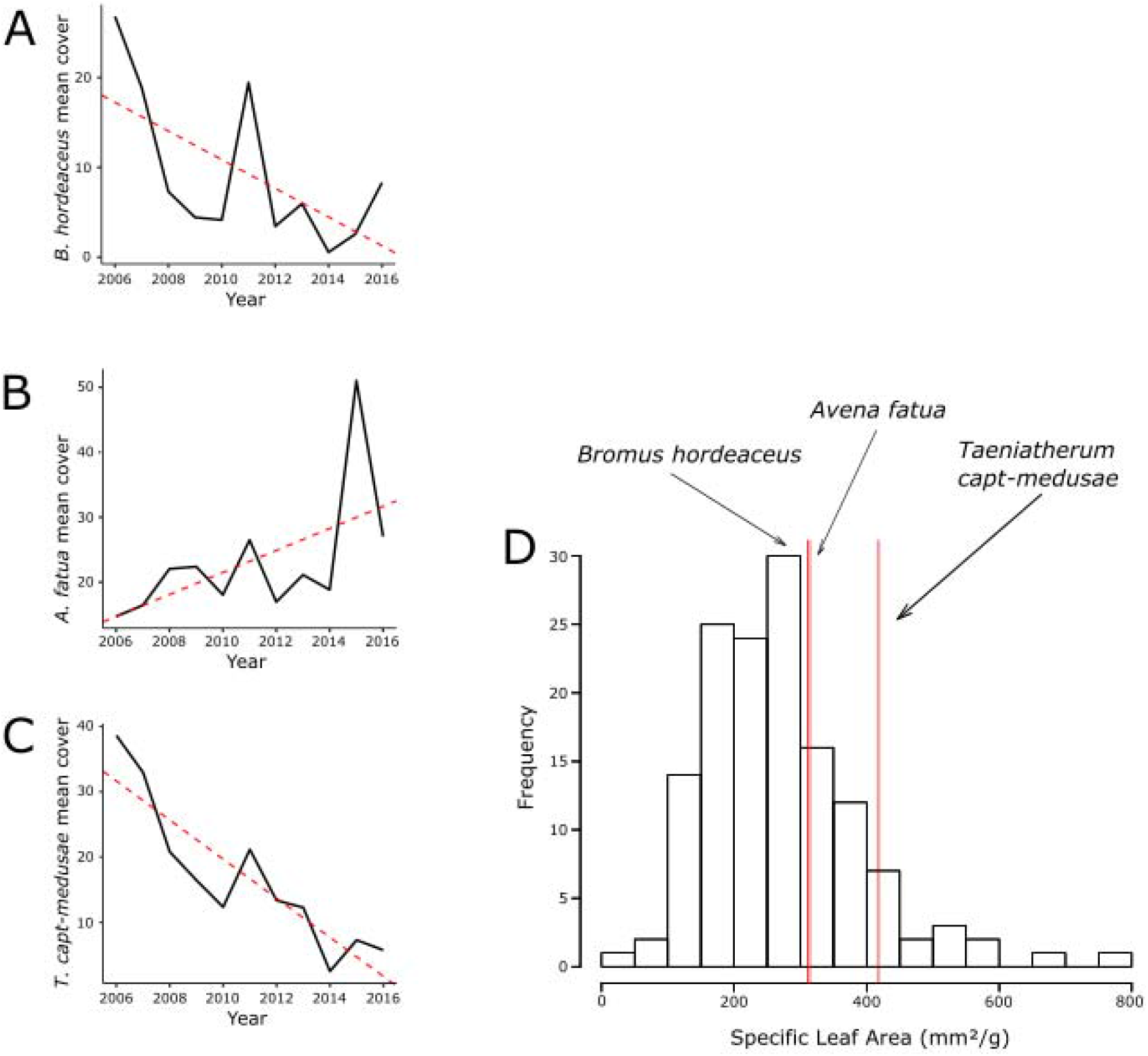
**A-C**: Changes in cover from 2006 to 2016 for the three grasses which best predict changes in nonserpentine community composition over time (**A**: *Bromus hordeaceus*; **B**: *Avena fatua;* **C**: *Taeniatherum caput-medusae*). **D**: Histogram of specific leaf area values for all nonserpentine species. Red lines indicate SLA values for *B. hordeaceus, A. fatua*, and *T. caput-medusae*.

